# Dynamic neural reconstructions of attended object location and features using EEG

**DOI:** 10.1101/2022.04.26.489558

**Authors:** Jiageng Chen, Julie D. Golomb

## Abstract

Attention allows us to select relevant and ignore irrelevant information from our complex environments. What happens when attention shifts from one item to another? To answer this question, it is critical to have tools that accurately recover neural representations of both feature and location information with high temporal resolution. In the current study, we used human electroencephalography (EEG) and machine learning to explore how neural representations of object features and locations update across dynamic shifts of attention. We demonstrate that EEG can be used to create simultaneous timecourses of neural representations of attended features (timepoint-by-timepoint inverted encoding model reconstructions) and attended location (timepoint-by-timepoint decoding) during both stable periods and across dynamic shifts of attention. Each trial presented two oriented gratings that flickered at the same frequency but had different orientations; participants were cued to attend one of them, and on half of trials received a shift cue mid-trial. We trained models on a stable period from Hold attention trials, and then reconstructed/decoded the attended orientation/location at each timepoint on Shift attention trials. Our results showed that both feature reconstruction and location decoding dynamically track the shift of attention, and that there may be timepoints during the shifting of attention when (1) feature and location representations become uncoupled, and (2) both the previously-attended and currently-attended orientations are represented with roughly equal strength. The results offer insight into our understanding of attentional shifts, and the noninvasive techniques developed in the current study lend themselves well to a wide variety of future applications.

**Open Practice Statement:** The data and analysis code will be made publicly available on the Open Science Framework (link to be updated upon publication).

**New & Noteworthy:** We used human EEG and machine learning to reconstruct neural response profiles during dynamic shifts of attention. Specifically, we demonstrated that we could simultaneously read out both location and feature information from an attended item in a multi-stimulus display. Moreover, we examined how that readout evolves over time during the dynamic process of attentional shifts. These results provide insight into our understanding of attention, and this technique carries substantial potential for versatile extensions and applications.

## Introduction

The visual environment contains so much information, and given that we have limited cognitive resources, visual attention plays an essential role to select the important information (Carrasco, 2011; Chun et al., 2011; Desimone & Duncan, 1995). Spatial attention is one way we can focus on the most relevant objects and locations for behavior, and filter out the irrelevant information. Spatial attention can accelerate target information accrual across eccentricity (Carrasco et al., 2006), and may speed the transition between sensory input and the formation of object representations (Di Russo et al., 2003; Foster et al., 2021; Hillyard & Anllo-Vento, 1998). In our daily lives, however, spatial attention is rarely static: when there are multiple objects or locations of interest, we may shift spatial attention frequently between them. Numerous studies have investigated the neural mechanisms of shifts of attention using various neuroscience tools (see reviews, Chica et al., 2013; Corbetta & Shulman, 2002; Miller & Buschman, 2013), and exploring the behavioral consequences of shifts of attention has become an important topic in the cognitive psychology literature (Carrasco, 2011; Dowd & Golomb, 2019; Egly et al., 1994; Folk et al., 2002; Paffen & van der Stigchel, 2010).

At the whole-brain network level, neuroimaging studies have established two separate fronto-parietal systems involved in different attentional operations: the dorsal attention network which is related to top-down goal-directed attention, responsible for the voluntary deployment of attention to stay focused on current goals, and the ventral attention network which is related to bottom-up, stimulus-driven attention, responsible for the reorientation to the salient or unexpected events in the environment (Corbetta & Shulman, 2002; Vossel et al., 2014). Neurophysiological evidence is consistent with top-down and bottom-up attention signals in frontal and parietal cortices (Buschman & Miller, 2007), and it has been well documented that fronto-parietal activation is associated with the control of orienting (Hopfinger et al., 2000; Kastner et al., 1999; Kelley et al., 2008; Peelen et al., 2004; Rosen et al., 1999; Yantis et al., 2002). Specifically, the superior parietal lobule (SPL) and medial regions of the prefrontal cortex show transient increases in neural activity when attention is disengaged from fixation and shifts to new peripheral locations (Kelley et al., 2008; Yantis et al., 2002). SPL is also shown to engage in covert shifts of attention between spatial locations (Gmeindl et al., 2016; Greenberg et al., 2010; Kelley et al., 2008, 2008; Zhang & Golomb, 2021), features (Greenberg et al., 2010), objects (Serences, 2004) and visual/auditory modalities (Shomstein & Yantis, 2004). Human EEG studies have further identified certain ERP components linked to spatial shifts of attention (Hillyard & Anllo-Vento, 1998; Kiss et al., 2008; Nobre et al., 2000; Yamaguchi et al., 1994), thought to be localized to extrastriate and parietal cortices (Di Russo et al., 2003; Hopf et al., 2000). Other studies have focused on neural timecourses of attentional shifts using electrophysiological signatures of EEG and single-unit recording (Khayat et al., 2006; Müller et al., 1998).

At the same time, human behavioral studies have revealed behavioral costs associated with shifts of attention. For example, reaction times are slower when attention must be shifted to a new location to perform a task, rather than holding attention at the same location (Maljkovic & Nakayama, 1996; Posner et al., 1980). Similar behavioral costs are found when a distracting stimulus captures attention away from a target location (Theeuwes, 1992). Furthermore, more recent studies have revealed that these dynamic shifts of attention bring additional challenges to our visual system to correctly bind location and features (Chen et al., 2019; Dowd & Golomb, 2019; Golomb, 2015; Golomb et al., 2014). Identifying visual objects requires our brain process both location and feature information (Holcombe, 2009; Reynolds & Desimone, 1999; Riesenhuber & Poggio, 1999; Singer, 1999; Treisman, 1996; Treisman & Gelade, 1980; von der Malsburg, 1999; Wolfe & Cave, 1999), and a common theory of feature integration suggests that attention serves as a glue to bind objects’ features together (Kristjánsson & Egeth, 2020; Nissen, 1985; Treisman & Gelade, 1980). During rapid shifts of attention -- and when spatial attention is otherwise disrupted or spread across different locations -- different types of feature binding errors can occur (Chen et al., 2019; Dowd & Golomb, 2019; Golomb, 2015; Golomb et al., 2014; Jones et al., 2021).

To study dynamic shifts of attention and understand how these behavioral consequences link to shifts of attention at a neural level, it is essential to have tools that can accurately recover neural representations of both feature and location information, and do so across a shift of attention with high temporal resolution. On the rise of machine learning and multivariate pattern analyses in recent years, many fMRI studies have made efforts to decode or reconstruct location and/or feature selective responses in the human visual cortex (see De Martino et al., 2008; Naselaris et al., 2011; Norman et al., 2006 for reviews). By making prior assumptions of organization of feature space, encoding models have advantages to reconstruct population-level response profiles of the sensory cortex (Sprague & Serences, 2015). The Inverted Encoding Model (IEM), one example of an advanced encoding model of neural representation, has been successfully utilized to reconstruct location or feature selective response profiles in both visual perception and visual working memory (Brouwer & Heeger, 2009, 2011; Foster et al., 2017; Scolari et al., 2012; Sprague & Serences, 2013).

Despite these recent advances, fMRI has inherently poor temporal resolution because of the lag of hemodynamic response. This makes fMRI a suboptimal tool to study the dynamic process of neural representations across attention shifts. Electroencephalography (EEG) and magnetoencephalography (MEG) have millisecond-level temporal resolution and make better candidates to reveal the dynamics of neural information processing. Previous studies have found EEG and IEM could be exploited to reconstruct visual perceptual information and working memory content (Foster et al., 2016, 2017, 2021; Garcia et al., 2013), but to our knowledge this has never been attempted across dynamic shifts of attention.

In the current study, we used EEG and IEM to reconstruct the neural response profiles during dynamic shifts of attention. Our design has two unique advances over prior studies. First, we focus on simultaneous readout of location and feature information from an attended stimulus, and how that readout evolves over time. Second, we used a multi-stimulus design, where two stimuli were presented but only one was attended at any given moment. In most previous studies, only one stimulus was presented, and the algorithm was run to reconstruct its location or feature (e.g. Foster et al., 2016). In that case the decoded information could come from two sources: the signal could be directly driven by the sensory information, and/or by the *attended* information. Therefore, to better understand shifts of attention and recover the content of attended information specifically, we presented two stimuli simultaneously and deliberately maintained the same visual information while manipulating spatial attention.

Prior studies have used EEG steady-state visual evoked potentials (SSVEPs) to access which of multiple items is being attended via frequency tagging, where each stimulus is tagged by presenting it repeatedly at a certain temporal frequency, which entrains the neural signal (Müller et al., 1998; Norcia et al., 2015). In the current study, however, we are interested in independently reconstructing *both* spatial and feature information. Thus, rather than using frequency-tagging, we presented the two stimuli at the same frequency, such that the generated SSVEP signal reflects both stimuli. We then conducted machine learning analyses to test whether we can reconstruct the *attended* location and feature information from this common signal. We are particularly interested in tracking how this recovered information updates with a shift of attention. Our goals were thus to establish: (1) whether this technique can produce reliable reconstructions of attended location and feature information from multi-stimulus displays; and (2) if we can track how these reconstructions change over time across dynamic shifts of attention.

## Methods

### Participants

25 subjects (8 male, 17 female; mean age = 21.56) participated in the experiment for monetary compensation ($15/hour). All participants reported having normal color vision and normal or corrected-to-normal visual acuity. Three additional participants were excluded due to poor behavioral performance (change detection accuracy in Hold trials < 10%; the rest of participants >70%, see *Stimuli and Procedure*). All participants provided written informed consent, and study protocols were approved by The Ohio State University Behavioral and Social Sciences Institutional Review Board.

### Behavioral Task

The stimuli consisted of one black fixation cross and two colored, flickering, square-waved gratings presented on a solid gray background with luminance of 37.5 cd/m2. The size of the fixation cross was 1° and displayed at 2° visual angle below the center of the screen. The size of each grating was 8° visual angle in diameter and displayed at 2° above and 6° left or right of the screen center (Figure 1). The spatial frequency of the gratings was 4 cycle/dva. The orientations were chosen from a set of 9 orientations (0°, 20°, 40°, 60°, 80°, 100°, 120°, 140°, 160°), such that the two gratings displayed were always 60 degrees apart (clockwise or counterclockwise), resulting 18 different stimulus pair combinations. Each orientation was then added an independent jitter ranging from −5° to 5°.

**Figure 1.**
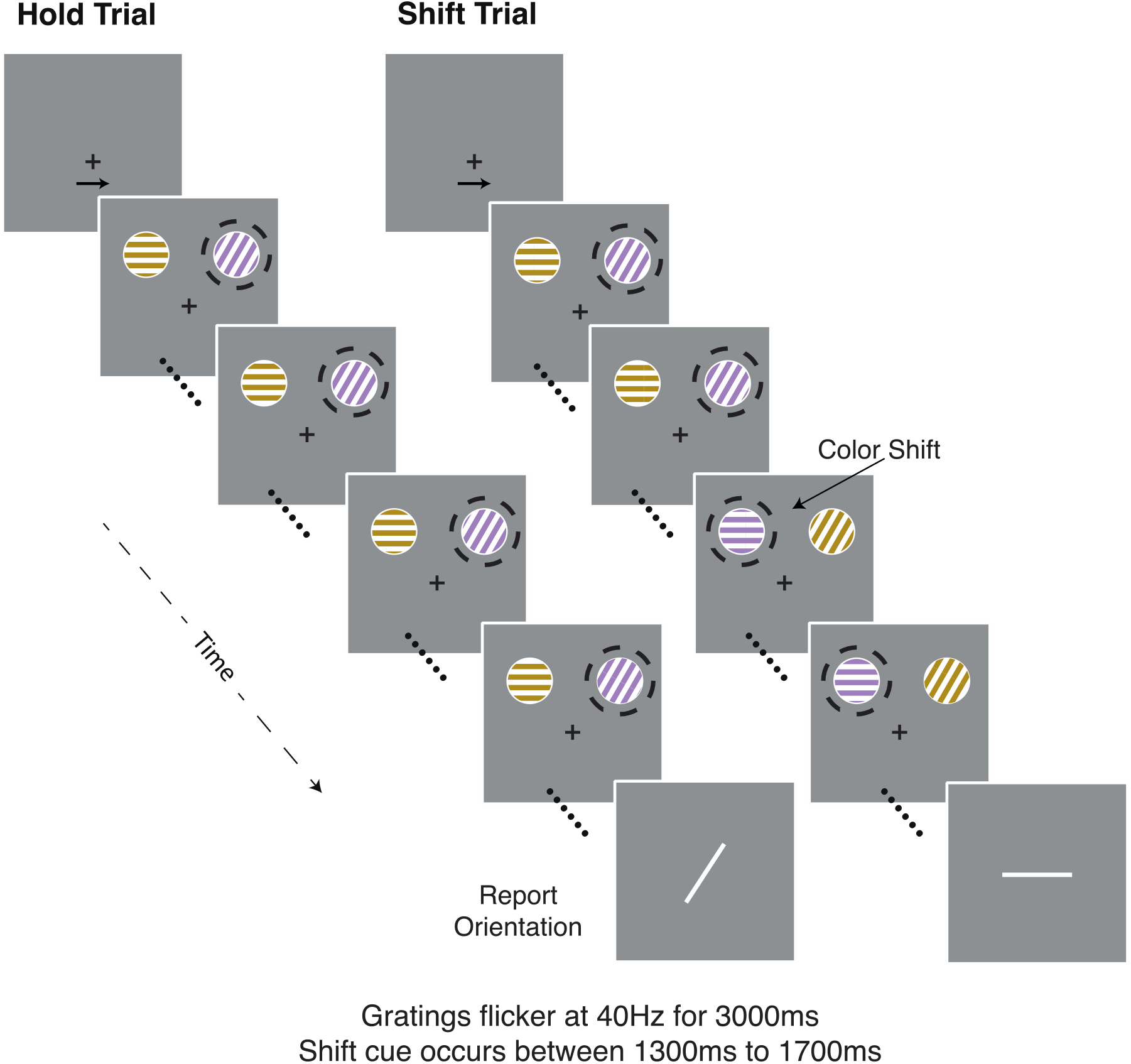
Example trial sequences for Hold and Shift Attention trials. Example here shows sequences for a participant asked to covertly attend the purple grating (half of the participants attended the gold grating instead). Dashed circles (not actually shown to participants) indicate the to-be-attended item over time. On Switch trials (randomly intermixed with Hold trials), the colors of the gratings switched in the middle of the trial and participants had to shift attention to track the purple (or gold) one. Participants were instructed to monitor the attended item for subtle orientation changes and press a button when one was detected. At the end of trial, participants were asked to rotate an orientation bar to match the orientation of the most recently attended grating.

One grating was colored purple (L=70, a*=28.4, b*=-21.4), and the other one was colored gold (L=70, a*=11.6, b*=97.4). In order to generate SSVEPs, the contrast of the gratings was reversed (e.g. purple to white to purple) at 40Hz. Participants were asked to always covertly attend to either the purple grating or the gold grating (color balanced across participants), all while keeping their eyes fixated on the fixation cross. The to-be-attended color was determined on a participant-wise basis: 13 participants always attended the purple grating during their session and 12 always attended the gold grating. The purple and gold gratings were equally likely to appear in the left or right positions at the start of the trial, and participants were instructed to covertly shift their attention if the colors switched positions (described below).

Before each trial, participants were shown a screen with a fixation cross and a black arrow above it that pointed left or right, indicating where the to-be-attended target grating would appear at the beginning of the trial. We included this additional spatial cue to avoid visual search and/or attention shift effects at the beginning of the trial. When they were ready to begin the trial, participants pressed the space bar. The two colored, flickering gratings appeared on the screen and were displayed for 3000ms.

There were two spatial attention conditions: In half of the trials, the colors of the two gratings remained the same throughout the trial, so participants attended to the same item/location the entire trial (“Hold condition”). In the other half of trials, the two gratings switched colors midway through the trial (i.e., the purple grating turned gold, and the gold grating turned purple). Once the two gratings switched their colors, participants needed to immediately shift their spatial attention to the other grating (“Shift condition”). On Shift trials the two gratings swapped colors but preserved their original orientations, so the spatial shift resulted in attending a new grating whose orientation was 60° different from the original one. Hold and Shift trials were intermixed and randomized in each block, such that participants could not predict whether a shift would take place at the beginning of the trial. The onset of the shift cue was randomly picked for each trial from a uniform distribution ranging from 1300ms to 1700ms after the stimulus onset.

To confirm that participants maintained their attention on the correct grating, each grating had 0, 1, or 2 subtle orientation changes (10°) during the trial. Participants were instructed to immediately press the “s” key when they detected an orientation change in the attended grating. They were also told to disregard any changes in the non-attended grating. Particularly, if the current trial was a shift trial, once the color-switch happened, participants needed to monitor and report the subtle orientation change in the newly attended grating and ignore the previously attended grating. At the end of trial, participants were also asked to rotate an orientation bar (appearing on the screen center) to match the orientation of the most recently attended grating and press the spacebar to confirm their answer.

To confirm that participants maintained fixation on the fixation cross while covertly attending the grating, we performed gaze-contingent eye-tracking. If a participant’s eye position deviated more than 1.5 dva from the fixation cross during the period while the flickering gratings appeared on the screen, the trial was aborted immediately and repeated at a random time later in the block.

The study was scheduled in two sessions. In the first session, participants completed two blocks of the main behavioral task without EEG, to familiarize themselves with the task. The second session (scheduled at a later time) was the official EEG session. During this 2-hour session, participants completed up to 12 blocks of the task (each containing 48 trials; 24 per condition) while EEG data were collected. Participants who completed at least 10 blocks (480 trials) were included in the analyses (M = 11.72 blocks).

### Experimental Setup

All stimuli were presented using MATLAB (MathWorks, Natick, MA) and the Psychophysics Toolbox (Brainard, 1997; Kleiner et al., 2007; Pelli, 1997) on an Apple Mac Mini. Participants were seated 80cm away from a 27-in. CRT monitor with a resolution of 1280*1024; and a refresh rate of 120 Hz. Participants’ eye position was monitored and recorded using an Eyelink 1000 system (SR Research, Ontario, Canada). A chin rest was used to stabilize participants’ head position. The CRT monitor was color calibrated with a Minolta CS-100 (Minolta, Osaka, Japan) colorimeter.

Scalp EEG activity was recorded while subjects performed the behavioral task in a shielded testing room. Each subject was fitted with an elastic cap containing 64 active Ag/AgCl electrodes arranged in an extended 10-20 layout, recorded via a BrainProducts actiCHamp Amplifier at a sampling rate of 1000Hz. Two additional electrodes (TP9, TP10) were attached to the left and right mastoids via electrode stickers. Electrode impedances were reduced to <25 kΩ before the commencement of each experiment session.

### EEG preprocessing

EEG data preprocessing was done using EEGLAB (Delorme & Makeig, 2004) and custom MATLAB scripts. We first downsampled the EEG data to 250Hz and re-referenced to the mean activity of all electrodes offline. Then we applied a band-pass filter from 0.1 to 58 Hz (using “pop_eegfilternew.m” in EEGLAB). The data were segmented into epochs corresponding to each trial, by taking EEG activity for each electrode from −500 ms to 3500ms relative to the start of that trial. (The time period when the stimuli were presented on the screen was 0ms to 3000ms.) We removed epochs in which the peak-to-peak range of any electrode was larger than 50 muV during the stimulus display (from 0ms to 3000ms relative to the start of each trial). Each epoch was then visually inspected to confirm no further artifacts. On average, 12.81% of trials (SD: 2.52%) were discarded for each participant after the preprocessing.

Our experimental design (described below) perfectly balanced trial counts across conditions, but after excluding noisy trials, the counts may not be fully balanced within each participant. Because an imbalance in the initially attended location (left vs right) could influence training of the models, we re-balanced the attended target location to equate the number of trials on which target was on the left side of screen in the beginning or on the right side of screen by randomly selecting a subset of trials from the larger group. Because each random selection caused a small number of trials to not be included in the final analyses, we repeated the selection process 100 times and applied all analyses for each selected dataset. We report the final averaged results to minimize the random selection effects.

### Behavioral Analyses

#### Change Detection Task

We calculated the d-prime for the change detection task. Each trial may have zero, one, or two orientation changes. Because two orientation changes in one trial could be displayed very close to each other, participants may respond by pressing the response key longer but not pressing twice. Because this change detection task was primarily intended to encourage and verify participant compliance, to simplify our analyses, we combined trials with one and two orientation changes. *Hit* trials were defined as trials where participants successfully detected any changes when there was at least one change. False Alarm trials were defined as participants reporting one or two changes when the trial had zero changes. d’ was calculated as:

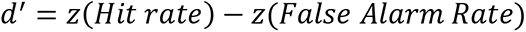

To avoid infinite values, we manually defined the minimum and maximum probability of each rate as 1/N and (N-1)/N, where N is the number of trials of that condition.

#### Post-trial Orientation Report

For the post-trial continuous report task, the difference between the correct orientation and the reported orientation was calculated as the “report error” for each trial. The report error range is from −90° to 90°. We realigned the direction of report error in the shift condition so that a positively signed report error means the reported orientation was attracted towards the orientation of the initially attended item (+60°); a negatively-signed report error means the reported orientation was repulsed away from the initially attended item’s orientation. On hold trials, the report error was mock aligned to match the shift condition (and eliminate any systematic clockwise/counterclockwise bias). We then fit the distribution of report error with a probabilistic mixture model (Bays et al., 2009; Zhang & Luck, 2008). The model assumes the distribution of report error comes from two sources (Formula 1): one von Mises distribution (ϕ) accounting for the probability to correctly report the target orientation, with a flexible mean (μ) allowing the model to capture any systematic bias from the target orientation and a flexible concentration parameter (κ) to capture precision; and one uniform distribution accounting for the probability (γ) of random guessing. Note: because we did not observe any large “swap” errors (see Figure 3), we chose this simpler mixture model without a swap error distribution.

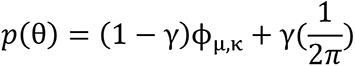

**Figure 2.**
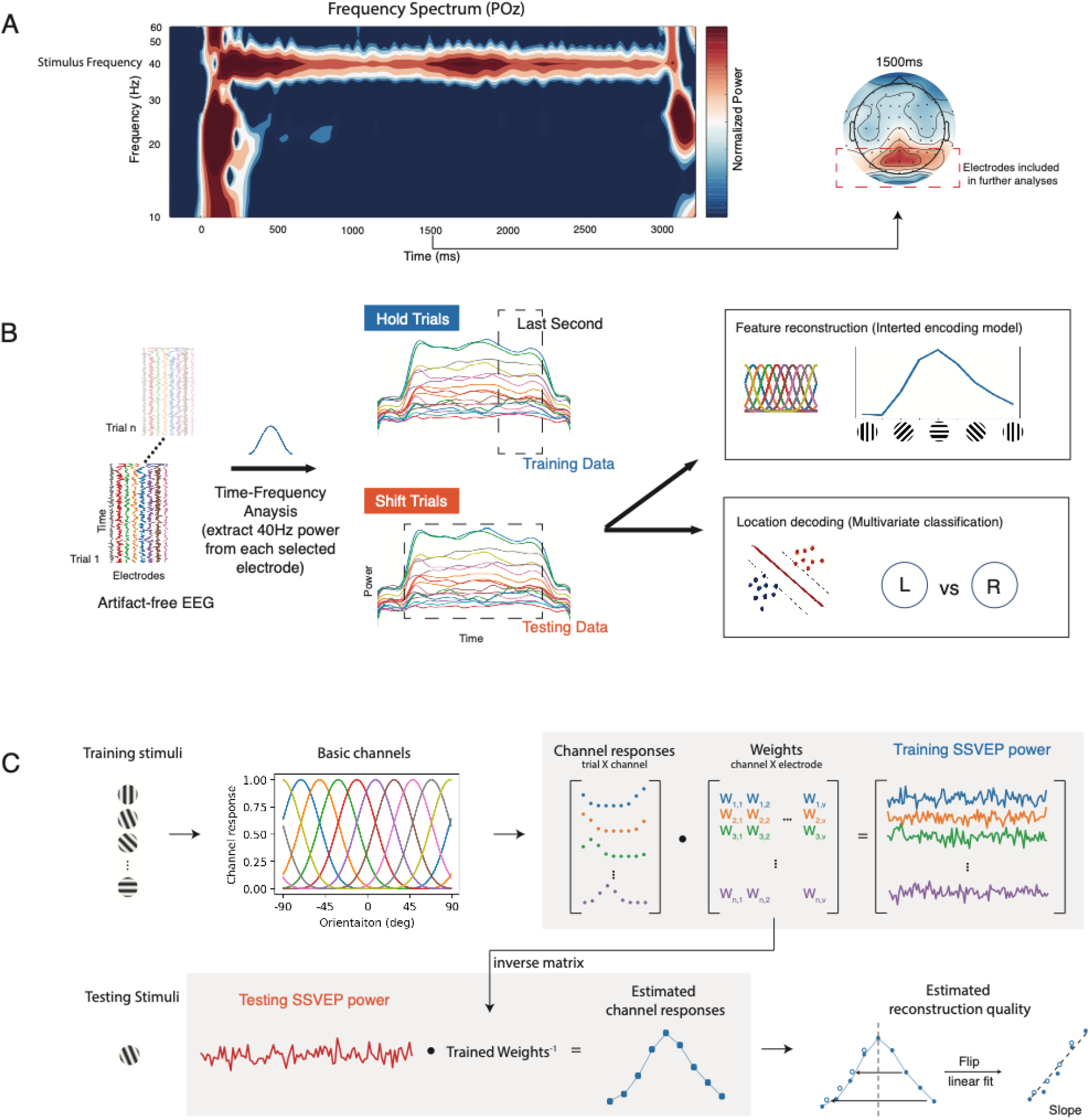
Overview of EEG analysis procedure. A) Time-frequency spectrum of example electrode (POz), showing the increased power in the 40Hz stimulus frequency band. Scalp distribution shows SSVEP power in the 40Hz band was strongest among parieto-occipital electrodes, as expected. B) Overview of EEG analysis pipeline for reconstructing attended feature and spatial information. C) Schematic for the feature reconstruction (Inverted Encoding Model) process. See Methods text for details.

**Figure 3.**
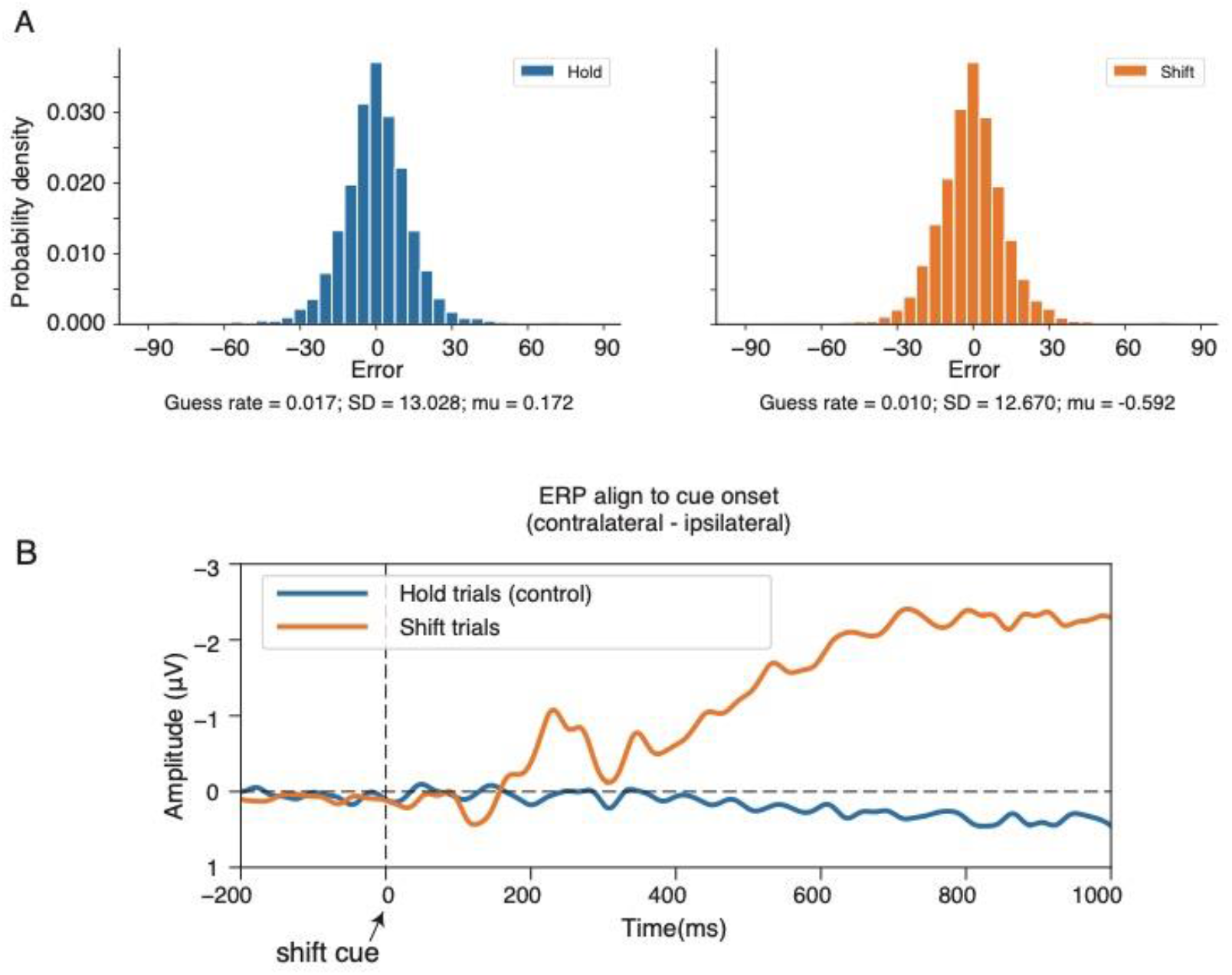
Behavioral and ERP results. A) Behavioral orientation report error distributions for hold and shift trials. We aligned the directions of the errors for shift trials so that the initially attended orientation was always represented at +60°. Text below plots gives the parameters from the mixture-modeling analysis: guess rate = probability of random guessing, SD = standard deviation of target distribution, and mu = mean shift of target distribution; values are averaged across subjects. B) ERPs from electrodes P7/P8, PO7/PO8, P3/P4 and O1/O2 aligned to shift cue onset time (for shift trials), calculated for trials with contralateral vs ipsilateral attended location. Because there were no actual shift cues in hold trials, we randomly selected time points during the shift cue window to align the hold trials (control data) accordingly.

For each participant and each condition, we fit the model by applying Markov chain Monte Carlo using MemToolbox (Suchow et al., 2013). The best-fitting parameters (maximum likelihood estimate) were compared between conditions. We also tested whether there were feature distortions in each condition by comparing the mean shift parameter (μ) to zero. We additionally calculated the mean signed error (without mixture modeling) for each participant and each condition as a non-modeling measure to determine whether the mean of the report error distribution for each condition was significantly shifted from zero.

### Manipulation Check: Event-related potentials (ERPs) Analysis

As another way of confirming participants were correctly allocating attention to the target orientation, especially on shift trials, we analyzed ERP data aligned to the shift cue (in the hold trials, we randomly picked a time point at each trial as the mock “shift cue” time). We averaged the signal amplitude from a subset of posterior and parietal channels (P7/P8, PO7/PO8, P3/P4 and O1/O2) based on the previous literature (Hakim et al., 2019), and subtracted the baseline EEG activity from 400ms to 0ms before the shift cue to calculate the ERPs. We sorted trials based on the attended side and calculated the difference waveforms by subtracting signals from contralateral side to ipsilateral side. We hypothesized that if attention was correctly shifted to the new target when the shift cue appeared, we should observe a robust N2pc component on shift trials, but not hold trials (Kiss et al., 2008). We calculated the mean N2pc amplitude by averaging the difference signals from 200ms to 300ms. We also calculated the contralateral delay activity (CDA) by averaging amplitude from 400ms post-shift cue onset to the end of the trial.

### Main EEG Analyses: Pipeline for reconstructing attended spatial and feature information

#### Time-frequency analysis

Our main analyses rely on time-frequency analyses of the preprocessed EEG signal. Below we describe the steps to extract the SSVEP power over time, which is then used for decoding attended location (see *Multivariate classification*) and attended orientation (see *Inverted Encoding Model*). This pipeline is visually depicted in Figure 2. Because our main emphasis is on reconstructing attended feature and location information across shifts of attention, we use the Hold trials as training data and the Shift trials as the testing data for the models.

First, to validate that our design evoked significant SSVEPs, we calculated the EEG power spectrum. Figure 2A shows an example electrode channel (POz) illustrating the increased power in the 40Hz frequency band (the stimulus frequency), with the spatial topography of the SSVEP signal maximal over the parietal-occipital electrodes.

To extract the timepoint-by-timepoint SSVEP power for the main analyses, we first applied a frequency-domain Gaussian shaped filter to the epoched artifact-free EEG signal for each trial (Cohen & Gulbinaite, 2017). The analysis is done using custom Python and Matlab scripts. A Fourier transform was applied to the padded signal to convert it from time-domain to frequency-domain. The frequency-domain EEG signal was point-wise multiplied by a gaussian shaped filter with peak frequency at 40Hz and full-width at half-maximum (FWHM) at 3 Hz. An inversed Fourier transform was then applied to recover the time-domain EEG signal. Finally, to extract the instantaneous power value of SSVEP, we applied a Hilbert transform to the filtered EEG data. To better deal with the edge effect, the signal was padded with 500ms blank data in both ends before the time frequency analysis. The padded data were removed after the analyses to maintain the same length as the original signal. To maximize our temporal resolution, we tested different wavelets with FWHM ranges from 0.5Hz to 5Hz and found at least 3Hz was required to achieve a reliable orientation reconstruction.

The above analysis results in a *m*n*t* matrix for each participant and each condition representing the spatiotemporal pattern of SSVEP power, where *m* is the number of electrodes, *n* is the number of trials, and *t* is the number of time points. To avoid overfit and reduce computational demands, the 17 posterior channels (P7, P5, P3, P1, Pz, P2, P4, P6, P8, PO7, PO3, POz, PO4, PO8, O1, Oz, O2) were selected for input to the decoding and encoding model, as previous literature has reported that SSVEP is most commonly observed in these posterior electrodes (Norcia et al., 2015; also see Figure 2A).

#### Inverted encoding models (Reconstructing attended orientation)

To reconstruct attended feature information on Shift trials, we used a cross-condition training and test routine. We trained the model based on the patterns of EEG SSVEP power and orientation of the attended items on *Hold* trials, and then inverted the model weights to reconstruct the attended orientation on *Shift* trials. We first applied this inverted encoding model (IEM) procedure to the stable attention periods of the shift trials, defining the before-shift period as the first second following stimulus onset (time 0 to time 1sec), and the after-shift period as the last second prior to stimulus offset (time 2sec to 3sec). SSVEP power was averaged over each time window for each electrode and participant. For both stable attention periods – as well as the dynamic reconstruction analyses below – we used identical training data, selecting the second window (time 2sec to 3sec) of Hold trials as the training dataset. The use of common training data ensures any differences in test results are not due to the differences in the training data, and we selected the last second of hold trials (rather than the first second) because there was no uncertainty at that point in the trial as to whether or not a shift would occur, so this period was the most pure hold-attention period.

For the dynamic reconstruction of attended feature over time analysis, we trained the model on the same final-second time window of Hold trials, and then tested the model on the timepoint-by-timepoint Shift data. We alternatively considered using a model with separate training data for each timepoint (train Hold time(t), test Shift time(t)), but there are both theoretical and practical advantages of using common training data for each reconstruction (Sprague et al., 2019). (Preliminary analyses using the timepoint-by-timepoint train and test procedure gave us similar, though noisier results.)

For the IEMs, we followed similar approaches as previous literature (Garcia et al., 2013; Sprague & Serences, 2015; Figure 2C). We assumed the signal at each electrode reflects the linear sum of 9 different hypothesized orientation channels (basis set). The response function of each basis channel is modeled as a half sinusoid raised to the 8^th^ power, where the centers of the 9 response functions are circularly distributed across feature space (20°, 40°, 60°, …, 180°). We repeated the process described below 19 times for each model, iteratively shifting the center of each response function 1° each time. Iterative shifting of basis sets allows for more accurate reconstructions across the full orientation space (Kok et al., 2013; Lorenc et al., 2018; Scotti et al., 2021).

The IEM model assumes a linear relationship between the EEG signal and channel tuning functions. During the training stage, a weights matrix is estimated as follows:

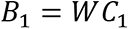

where B_1_ (m electrodes * n trials) is the observed EEG signal (SSVEP power) at each electrode in the training set, C_1_ (k channels * n trials) is the response function of the hypothesized orientation basis set channels, and W (m electrodes * k channels) is the weight matrix that characterizes a linear mapping from channel space to electrode space. The weight matrix W is derived via ordinary least-square estimations as:

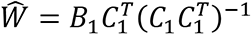

where *Ŵ* (m electrodes * k channels) is the least-square solution.

In the test stage, we inverted the model to transform the test data B_2_ (m electrodes * 1 trial) to the estimated channel response *̂*_2_ (k channels * 1 trial) using the estimated weight matrix *Ŵ* :

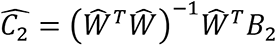

The output of the model is the estimated channel response for each test trial (and/or test timepoint). After iterative shifting, these channel-tuning functions (CTFs, Foster et al., 2017) were circularly shifted to align all trials to a common center for statistics and illustration purposes; for our figures the aligned reconstruction plots were centered on 30° (range −60° to 120°), with 0° indicating the orientation of the initially attended item and 60° the orientation of the second attended item.

Because in the hold condition the attended orientation stays the same and in the shift condition the attended orientation changes by 60° in the middle of the trial, if IEM correctly models the attended orientation, we should observe CTFs shift their peak center from the initially attended orientation (0°) to the newly attended orientation (60°).

To quantify the orientation sensitivity of the CTFs, we calculated linear slope as an index of orientation sensitivity (Foster et al., 2017; Samaha et al., 2016; van Moorselaar et al., 2018; Yu et al., 2020). We calculated symmetric slope by reversing the sign of positive orientation channels and collapsing their channel responses with the corresponding negative degrees. Then we fitted a linear regression to obtain the linear slope as the sensitivity measure. Higher slope indicates greater orientation sensitivity. For shift trials, we calculated slope in two ways: relative to the initial attended item’s orientation (*CTFslope-O1*), and relative to the second attended item’s orientation (*CTFslope-O2*). Reliable reconstructions of attended feature information should show a *CTFslope-O1* significantly greater than 0 in the first part of the shift trial and a *CTFslope-O2* significantly greater than 0 in the second part of the shift trial (see Statistics section below).

#### Multivariate classification (Decoding attended location)

For the attended *spatial* information analyses, support vector machine (SVM) was applied to determine whether the attended location (left vs right) could be decoded from the spatial distribution of SSVEP power over time. Analogous to above, we trained the SVM on the last second of hold trials, using SSVEP power and the correct attended location for each trial, and then tested at each timepoint on the shift trials to predict its attended location. Because there were only 2 possible locations to attend, the chance level of the prediction is 50% (left vs right). We used custom python code and “SVC” function from “sklearn” package, using a linear kernel and regularization parameter set to 1.0.

#### Statistical significance tests

To determine significant time points for the above analyses, we used cluster-based permutation tests to correct for multiple comparisons and identify clusters of time points when the CTF slopes were significantly larger than 0 (significant orientation reconstruction) and/or location decoding performance was significantly better than chance (Cohen, 2014; Maris & Oostenveld, 2007). For each analysis, we first did a one sample t-test to detect time points with CTF sensitivity greater than 0 (or location decoding accuracy greater than 0.5). We used .05 as the alpha threshold (t = 1.711, one-sided, df = 24) to identify clusters of adjacent points, and computed the sum of all the *t* values within each cluster. We then compared the sum of t-values against a null distribution empirically specified with the Monte Carlo randomization procedure. The null distribution is calculated by randomizing the training and test labels and repeating the IEM procedure (or multivariate classification procedure, in case of location decoding performance) 1,000 times. We followed the same procedure as described above to compute the sum of t-values for the largest cluster for each of the 1,000 iterations, resulting a null distribution with 1,000 sums of t-values. We compared the sum of t-values of the correctly labeled data with the 95^th^ percentile of the null distribution to determine whether the cluster was above chance (one-tail alpha rate = .05).

## Results

### Behavior and ERP analyses confirm participants successfully performed the attention task

Behavioral analyses of the change detection task indicated that participants were able to allocate and maintain their attention to the correct location. Participants detected the orientation changes in the attended item significantly better than chance level on both Hold trials (dprime = 2.258, t(24) = 8.988, p<0.001) and Shift trials (dprime = 1.269, t(24) = 6.562, p<0.001). Post-trial continuous orientation reports similarly showed that participants reported the target orientation rather accurately, with probabilistic mixture models outputting low guess rates and high precision (small standard deviation) for both hold and shift trials (Figure 3A).

Given prior behavioral reports of feature distortions when attention is split across two locations (Chen et al., 2019; Dowd & Golomb, 2019; Golomb, 2015; Golomb et al., 2014), we also measured feature distortions (target orientation report either biased towards or repulsed away from the other item’s orientation). There was no evidence for distortion in hold trials: mu was not significantly different from zero (t(24)=1.017, p=0.318). However, for shift trials, mu was slightly but significantly negative (t(24)=2.372, p=0.025), indicating participants’ post-trial orientation reports were shifted away from the initially attended orientation (*repulsion* effect). We also assessed this in a model-free analysis by analyzing the mean of the entire error distribution. For hold trials, we did not observe a response bias (M=0.140; t(24)=0.642, p=0.527). For shift trials, we found the mean of reporting error was numerically negative and marginally significant (M=-0.526; t(24)=-1.930, p=0.066), consistent with a weak response bias away from the initially attended orientation.

As a preliminary analysis and sanity check of the EEG data, we also analyzed ERPs, with data aligned to the shift cue. Shifts of spatial attention are associated with characteristic ERP components, particularly a contralateral N2pc at the posterior/occipital channels, typically peaking from 200ms to 250ms at P7/P8, PO7/PO8, P3/P4 and O1/O2 (Luck, 2012). We observed a robust N2pc on Shift trials (Figure 3B), peaking at 229ms after the shift cue (M=-0.937 μV, t(24)=3.318, p=0.003). On Hold trials, no N2pc was present, as expected. Another ERP marker of selective spatial attention and maintaining objects in working memory is the CDA (Vogel et al., 2005; Vogel & Machizawa, 2004), which is apparent in Figure 3B for Shift trials, peaking at 700ms after the shift cue (M=-2.285 μV, t(24)=8.892, p<0.001). Note a robust CDA on hold trials would have been visible if the data were aligned to the stimulus onset, but it is not visible in Figure 3B because these ERP plots were aligned and baseline-adjusted to the non-existent shift cue.

### Feature information can be reliably reconstructed from multi-stimulus displays

Having confirmed that our behavioral task was successful at manipulating selective attention and evoking covert shifts of attention, we turned to our first main goal: Can the EEG IEM model reliably reconstruct attended feature information from these multi-stimulus displays? In other words, before attempting to track how neural reconstructions might change dynamically around the time of a shift of attention, we first needed to confirm that we could reconstruct the orientation that was attended in the first half of the trial (before any shift cue), and the orientation attended in the second half of the trial (well after the shift). Note that based on prior work we expected that attended *location* would be reliably decoded from the EEG signal during these static attention periods, and indeed that was true: The average location decoding accuracy in the before-shift static period was 0.651 (SD 0.125), significantly above chance (chance level: 0.5; t(24)=5.906, p<0.001). In the after-shift static period, the location decoding accuracy was 0.708 (SD 0.121), also significantly above chance (t(24)=8.442, p<0.001).

Critically, we could also reconstruct attended *feature* information during the static attention periods before and after the shift cue (Figure 4). In the before-shift static period, the reconstruction revealed two peaks: a primary peak centered on the orientation of the initially to-be-attended item (the current target), as well as a smaller peak centered on the orientation of the other item in the display. Thus, our technique is capable of reconstructing the orientations of two different items in the display, and of differentiating which one is at the current focus of attention. In the second half of the trial, once attention had fully shifted to the other item, the feature reconstructions successfully reflected this update, with the reconstruction peaking at the orientation of the now-attended item. Interestingly, unlike in the first half of the trials, the orientation of the other, unattended item was *not* strongly reconstructed in the second half of the trial. This suggests that in the current task, participants may have initially distributed some attention to the other item in preparation for a potential attention shift to the new target, but once the shift was executed, they more effectively filtered out the other item. This finding links well with recent behavioral work finding anticipatory sampling of attention prior to expected attentional shifts (Jones et al., 2021) and with working memory studies showing successful dropping of items no longer relevant (Lewis-Peacock et al., 2018; Souza et al., 2014).

**Figure 4.**
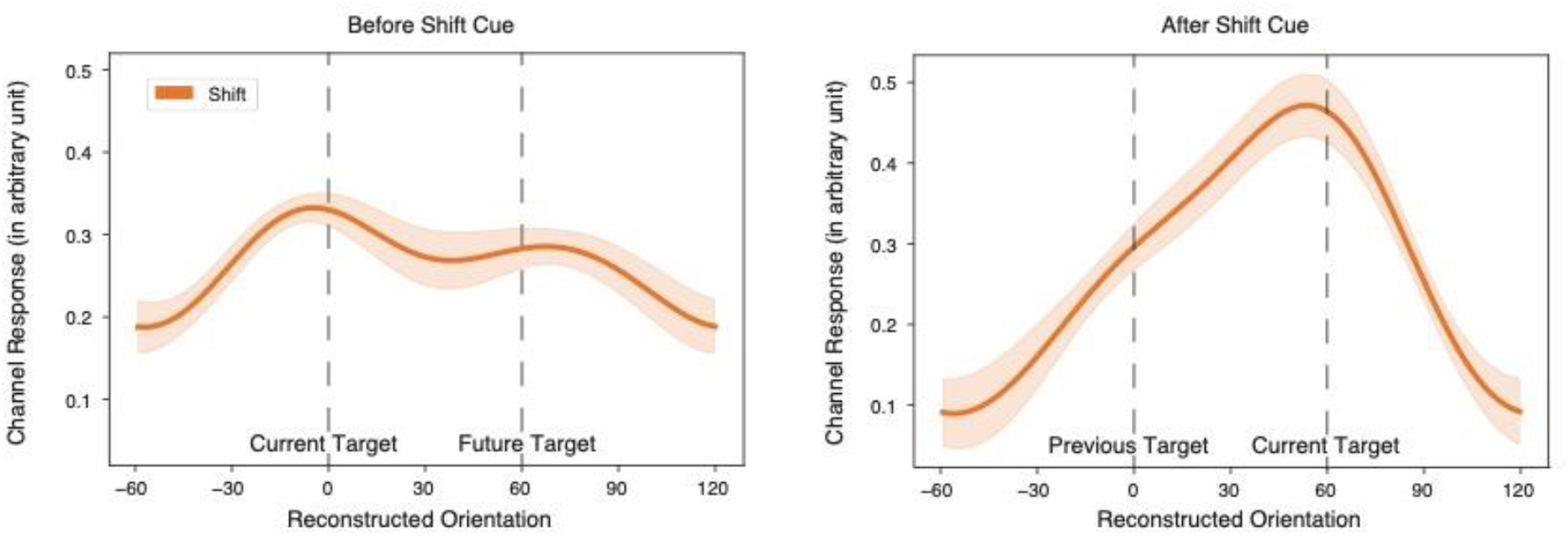
Orientation reconstructions during the static attention periods. IEMs were trained on Hold trials (final second of stimulation), and tested on Shift trials before and after the shift cue. The before-shift static period is the first second following stimulus onset (time 0 to time 1sec), and the after-shift static period is the last second prior to stimulus offset (time 2sec to 3sec). Before shift cue, the reconstruction showed two peaks: a strong reconstruction for the current target and a weaker reconstruction for the future target. After shift cue, there was primarily one stronger reconstruction for the current target.

### Reconstructed location and orientation both track the shift of attention

Next, we tested whether SSVEP and IEM reconstructions can dynamically track the attended orientation as spatial attention shifts to a different stimulus during the trial. Figure 5A shows the timecourse of feature reconstructions on shift trials, plotted as time-by-time channel tuning functions (CTFs) temporally aligned for each trial such that time 0 is the onset of the shift cue. Consistent with the static reconstructions, dynamic CTFs accurately reconstructed the orientation of the initially attended stimulus during the first half of the trial. Immediately following the shift, a period of poorer / ambiguous reconstruction was visible, followed by a settling of the reconstructions on the orientation of the newly attended stimulus (aligned at 60 degrees). Figure 5B plots these feature reconstructions another way, using CTF slope as a quantitative measure to assess reconstruction quality at each time point. We calculated CTF slope in two ways for each time point: centered on the orientation of the initially attended item (*CTFslope-O1)* and centered on the orientation of the newly attended item (*CTFslope-O2)*. During the first half of the trial, the orientation of the initially attended item was significantly reconstructed (*CTFslope-O1 >* 0, p<0.05 cluster-based permutation test) at all time points. After the shift cue there was a transient period (168-400ms post-cue) where both the initially and newly attended orientations were significantly reconstructed, and then eventually only the orientation of the newly attended item was recoverable. This indicates that our dynamic IEM approach successfully tracked the attended orientation(s) across the shift of attention. Moreover, the period where both orientations seemed to be represented – with overlapping timepoints where both CTFslope-O1 and CTFslope-O2 were significant – is particularly intriguing. Such a pattern is consistent with prior findings of temporal overlap in attentional facilitation during shifts of attention (Dowd & Golomb, 2019; Golomb, 2019; Khayat et al., 2006; Shulman et al., 1979), as we speculate on later in the Discussion.

**Figure 5.**
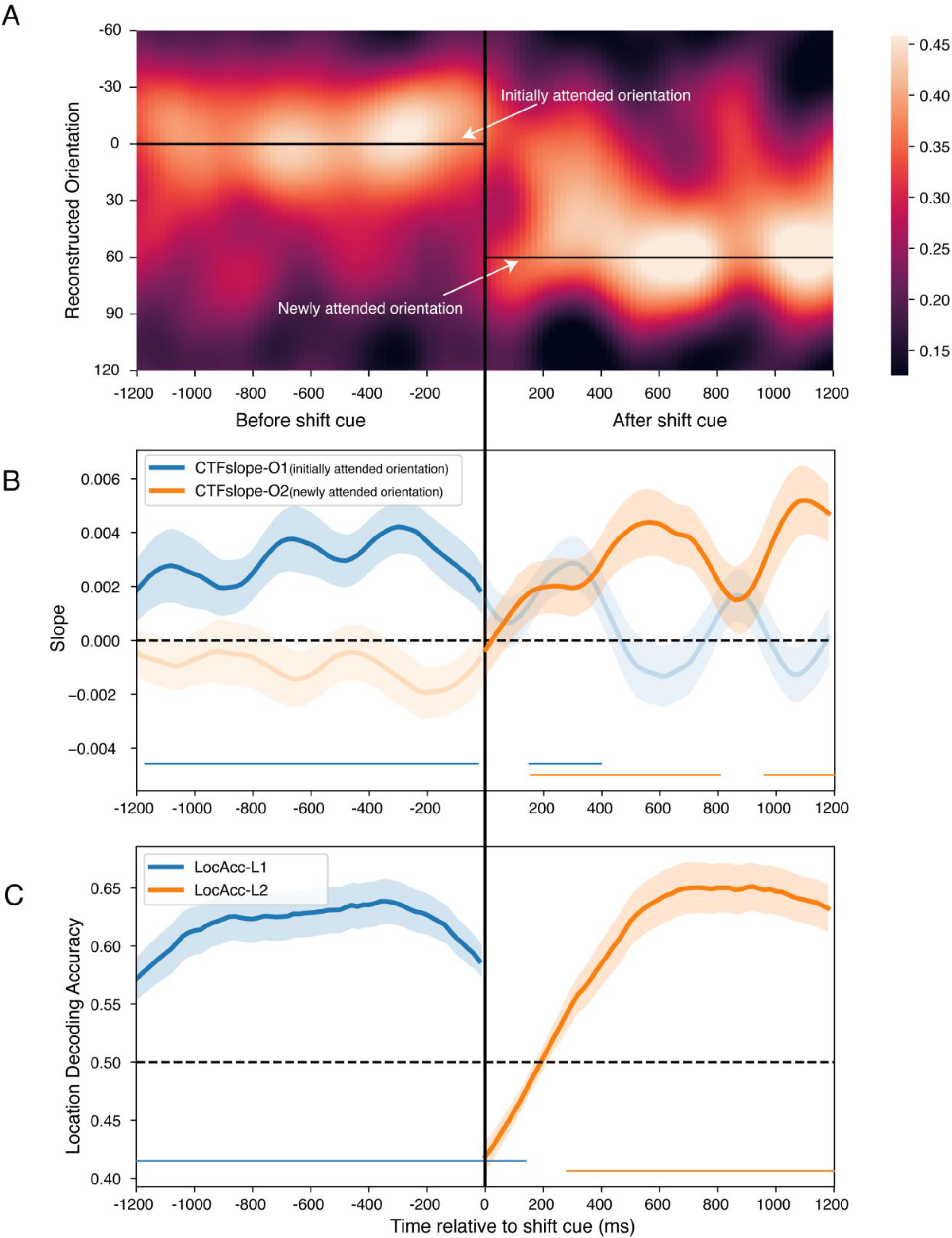
Feature reconstructions and Location decoding accuracy of Shift trials. A) Reconstructed channel tuning functions for each timepoint on Shift trials. Trials were aligned in time, such that time 0 was when the shift cue appeared, and in orientation space, such that the orientation of the initially attended item was centered at 0° and the newly attended item’s orientation was always represented at 60°. Colors reflect the amplitude of the reconstructed signal. B) Reconstruction slopes across time, calculated based on the initially attended orientation (*CTFslope-O1 = blue line)* and the newly attended orientation (*CTFslope-O2 = orange line*). C) Decoded location accuracy for each timepoint on Shift trials, aligned as in A-B. Before the shift cue, accuracy based on the correct initially attended location (*LocAcc-L1 plotted in blue for consistency with B)*; after the shift cue, accuracy based on the correct newly attended location (*LocAcc-L2 plotted in orange.* In B-C, the shaded error bars reflect ±1 SEM across participants, dashed black lines indicate chance level, and the solid bars along the bottom of the plots indicate timepoints that significantly differed from chance (cluster-based permutation tests).

We then examined the timecourse of attended location decoding (Figure 5C). This analysis was quantified as a simple decoding accuracy (attending left vs right location). Before the shift cue, we could decode the attended location (left vs right) consistently above chance (chance = .5; p<0.05, cluster-based permutation test). After the shift cue, the decoded location gradually shifted to the other side and became significantly above chance after 260ms, peaking around 600ms.

Cluster-based permutation tests were performed for both the dynamic feature reconstruction and dynamic location decoding analyses. Although it is important to keep in mind that these are quantified by different measures, and location is a two-way decoding while orientation is a continuous reconstruction, there are some intriguing comparisons between the two timecourses that may be interesting to speculate on. Comparing the attended feature and location timecourses revealed what could be characterized as multiple distinct periods: a stable pre-cue period, potentially three distinct transition stages, and a stable post-cue period. During the stable period before the shift cue, the correct currently attended location and orientation could both be significantly and robustly recovered from the EEG SSVEP signal. For the first 150ms following the shift cue, spatial attention appeared to still be primarily lingering at the initial location, though the signal was rapidly decaying to chance. During this time, neither orientation could be reconstructed above baseline. From timepoints 152-250ms post-cue, location decoding was not significantly different from chance, suggesting spatial attention was truly in transition. Strikingly, during this ambiguous spatial attention period, both the initial and the newly attended orientations could be significantly reconstructed. Since the location decoding analysis was simply a two-way decoder, we can’t resolve whether spatial attention was simultaneously at both locations or neither (or highly variable across trials), but we are clearly capturing a transitory period of ambiguous spatial attention, during which both items’ orientations were represented. Starting at 280ms post-shift, the location decoding became significant for the newly attended location; yet interestingly, both orientations could still be significantly reconstructed for another 100ms. Finally, starting around 400ms post-cue, only the correct newly attended orientation and location were significantly represented. Reconstruction slope and location decoding accuracy both continued to increase for another 100ms or so, plateauing into the post-shift stable period. As another interesting point of comparison, it is also potentially notable that the location decoding timecourses remained at a relatively constant and stable accuracy over the duration of the static-attention periods, whereas the orientation reconstruction slopes seemed to oscillate throughout the trial; one possibility is the feature reconstruction technique is more sensitive to oscillations of attention and/or divided attention, a speculation we revisit in the Discussion.

## Discussion

In the current study, we used EEG and IEM to reconstruct the neural response profiles during dynamic shifts of attention in high temporal resolution. Specifically, we demonstrated that we could simultaneously read out both location and feature information from an attended stimulus to produce reliable reconstructions of attended location and feature information from multi-stimulus displays. Moreover, we examined how that readout evolves over time during the dynamic process of attentional shifts.

Our study offers several methodological and theoretical contributions. In terms of methodological contributions, our study can be thought of as a proof of concept that EEG can be used to construct timecourses of the neural representations of attended features (timepoint-by-timepoint IEM reconstructions) and attended location (timepoint-by-timepoint decoding) during both stable periods and across dynamic spatial shifts of attention. Our approach builds off of prior studies using machine learning and IEM to decode/reconstruct the locations or features of a stimulus, either visually presented or in memory, from neuroimaging data (Brouwer & Heeger, 2009, 2011; De Martino et al., 2008; Foster et al., 2016, 2017, 2021; Garcia et al., 2013; Naselaris et al., 2011; Norman et al., 2006; Scolari et al., 2012; Sprague & Serences, 2013, 2015). However, unlike most of the previous studies, we focused on (1) both the location *and* feature information, (2) for an attended stimulus in a multi-stimulus display, (3) explored how the readout information evolved over time, and (4) showed that a model trained on Hold-attention trials could be used to reliably track the updating of neural representations on Shift-attention trials. To our knowledge this is the first study to successfully attempt this combination of goals. Of course, as we discuss more below, with all new approaches there is room for improvement and refinement, but the current results demonstrate that this approach is both feasible and carries substantial potential for versatile extensions and applications.

Moreover, in addition to the methodological contributions of this study, our results reveal some intriguing aspects of attentional updating that contribute to various theoretical issues in the attention literature. One aspect is how attended and unattended items are represented in a multi-item display. A number of prior studies have demonstrated that neural reconstructions of object features are more precise for attended than unattended items (Ester et al., 2016; Jehee et al., 2011), and that the attended orientation can be decoded from ambiguous stimuli (Kamitani & Tong, 2005). In the current study, we similarly found that we could reliably reconstruct the attended orientation during the static attention periods. Yet there was a notable asymmetry between the pre- and post-shift periods of the trial, with the results suggesting that participants were not exclusively attending to the initial item during the first half of the trial, but also allocating some attention to the other item in the display. In the first half of the trial, participants did not know if they would be holding attention or shifting attention, so there may have been some incentive to represent both items in the display, and then narrow the focus of attention in the second half of the trial. This pattern bears similarlities to the working memory literature, where studies have examined how representations change when items are added to or dropped from working memory (Balaban & Luria, 2017; Lewis-Peacock et al., 2018; Souza et al., 2014; Wan et al., 2020; Yu et al., 2020), except in the current study, only one orientation needed to be attended and held in WM at a time. We note that it is unlikely that participants were simply distributing their attention across both items in the first half, as the behavioral task required sustained focused attention on the attended item for the unpredictable and challenging change-detection task, and the results showed that participants were indeed focusing their attention on the current target item, as both the attended feature and attended location could be reliably extracted from the neural signal. Thus, it’s not that participants failed to perform the focused attention task, but rather the reconstructed signal was more contaminated by the non-target item before vs after the shift.

Another finding of the current study is that there appeared to be a transitional period following the shift cue during which both the previously attended and the currently attended orientations could be significantly reconstructed. In other words, after the spatial shift of attention, the previously relevant orientation was not immediately discarded, but was still temporarily represented in the neural signals. Because these reconstructions averaged across trials, it is difficult to say whether this effect was due to variable timing of attentional updating across trials or simultaneous representations of both items. However, a prior study in primate V1 found that during spatial shifts of attention, attentional enhancement is found for the item that is newly attended (distractor to target status) faster than attention is withdrawn from the initially attended item (target to distractor status) (Khayat et al., 2006). ERP evidence has also suggested that attention can be maintained at its previous location while it is simultaneously allocated to a new target object (Eimer & Grubert, 2014). Similar temporal overlap of attentional facilitation has been found when attention is updated across eye movements, resulting in a dual spotlight (Golomb, 2019) or soft handoff (Fabius et al., 2020; Marino & Mazer, 2018) of attention, and soft handoffs of attention are also found across hemispheres during multiple object tracking (Drew et al., 2014).

Indeed, the timecourse of attentional shifting has been debated over the years across behavioral (Duncan et al., 1994; Shulman et al., 1979; Wolfe, 1994), monkey neurophysiological (Khayat et al., 2006), and human EEG (Müller et al., 1998; Woodman & Luck, 1999) studies. Our current study offers a unique addition of providing several simultaneous measures that can track the timecourse of attentional shifts, including the N2pc, decoding of attended location, *and* reconstruction of attended orientation. The reconstruction timecourse for attended orientation revealed that the newly attended orientation first became significant around 168ms post-cue, similar to Khayat et al’s distractor-to-target latency of 144ms post-switch (Khayat et al., 2006). Meanwhile, the previously attended orientation was still significantly reconstructed at 400ms (substantially after Khayat et al’s target-to-distractor latency of 210ms). The location decoding timecourse crossed the chance point at 152ms and then became significant for the new location at 250ms. And the N2pc peaked 229ms after the shift cue. Meanwhile, both the location decoding and orientation reconstructions didn’t reach their peaks until 500-600ms after the cue. One takeaway from these data is that attentional shifting is perhaps better thought of as a more nuanced set of multiple processes or steps that unfold over an extended time window, rather than a single unitary switch.

Moreover, other previous literature has suggested that location plays a vital role in the process of binding features into cohesive objects (Treisman, 1996, 1998). When we talk about attention shifting from one object to another object, we generally do not separate the attended location and attended feature. But in fact, during the shift of attention, both the attended location and feature representations are updated, and location and feature representations may involve different brain regions (Ungerleider, 1994). An important question that this paradigm opens up is whether these two processes are temporally linked such that the timing of location updates is correlated with the timing of feature updates. The data presented here suggest some intriguing links, but an exciting future direction of this paradigm would be investigating correlations between the feature and location timecourses across subjects and/or trials. Because we only collected a single session of EEG data per subject, the current experiment was not powered to get reliable measures of transition timepoints in individual subjects, but future studies employing more extensive sampling may be better powered to investigate individual differences.

Another appealing direction for future applications of this technique would be to try to link individual or trial-wise behavior with the attended location and feature reconstruction measures. We did not find significant correlations between behavioral report measures and location or feature reconstructions in the current study, but we note that the behavioral tasks were not optimized to detect subtle variations in attentional state, but rather primarily meant to ensure that participants were attending to the correct item. As such, performance in the post-trial behavioral task (orientation report) was essentially at ceiling, exhibiting very low variability, and the frequency of the probes in the ongoing change detection task was too low to use for this purpose. That said, this paradigm may carry even more enticing potential for investigating attentional contexts that produce more behavioral errors and variability, such as divided attention (Dowd & Golomb, 2019), attentional capture by salient distractors (Chen et al., 2019), remapping across eye movements (Golomb et al., 2014), vigilance/distraction (Esterman et al., 2013; Rosenberg et al., 2015), and rhythmic oscillations of attention (Fiebelkorn et al., 2013; Landau & Fries, 2012).

One limitation of the current study is that the measures we used for assessing the attended feature and attended location representations are not directly comparable in terms of quality. We chose to focus on a single shift of attention between two fixed locations in the current study for a well-powered and clean proof of concept, and thus our attended location measure was limited to two-way decoding. In principle, future tasks could be designed such that a continuous reconstruction measure (IEM or other model-based technique) could be used to evaluate both attended location and attended orientation on the same scale, though likely multiple sessions of EEG data would be needed per subject.

In conclusion, by applying IEM and machine learning methods to EEG data, we simultaneously reconstructed feature representations and the location of spatial attention over the shift of attention in a multi-stimulus design. Our results showed that both feature reconstructions and location decoding dynamically track the shift of attention, and that there may be timepoints during the shifting of attention when (1) feature and location representations become uncoupled, and (2) both the previously-attended and currently-attended orientations are represented with roughly equal strength. The results offer insight into our understanding of attentional shifts, and the techniques developed in the current study lend themselves well to a wide variety of future applications.

## Acknowledgments

This work was supported by grants from the National Institutes of Health (R01-EY025648) and from the National Science Foundation (NSF 1848939) to JG. We thank Maurryce Starks for assistance with data collection and helpful discussion.

